# Distinct domains of Me31B interact with different eIF4E isoforms in the male germ line of *Drosophila melanogaster*

**DOI:** 10.1101/2021.03.23.436655

**Authors:** Carla Layana, Emiliano Salvador Vilardo, Gonzalo Corujo, Greco Hernández, Rolando Rivera-Pomar

**Affiliations:** Centro Regional de Estudios Genómicos, Facultad de Ciencias Exactas, Universidad Nacional de La Plata, Boulevard 120 N° 1459, 1900-La Plata; Translation and Cancer Laboratory. Unit of Biomedical Research on Cancer, National Institute of Cancer (Instituto Nacional de Cancerología, INCan). 22 San Fernando Ave., Tlalpan, 14080-Mexico City, Mexico; Centro de Bioinvestigaciones, Universidad Nacional del Noroeste de Buenos Aires, Av. Presidente Frondizi Km 4, 2700-Pergamino, Argentina

**Keywords:** Me31B, eIF4E, mRNP, translation initiation, P-bodies, *Drosophila*

## Abstract

Eukaryotic translation initiation factor 4E (eIF4E) is a key factor involved in different aspects of mRNA metabolism. *Drosophila melanogaster* genome encodes eight eIF4E isoforms, and the canonical isoform eIF4E-1 is a ubiquitous protein that plays a key role in mRNA translation. eIF4E-3 is specifically expressed in testis and controls translation during spermatogenesis. In eukaryotic cells, translational control and mRNA decay is highly regulated in different cytoplasmic ribonucleoprotein foci, which include the processing bodies (PBs). In this study, we show that *Drosophila* eIF4E-1 and eIF4E-3 occur in PBs where might play a role in mRNA storage and translational repression. We also demonstrate that the DEAD-box RNA helicase Me31B, a component of PBs, physically interacts with eIF4E-1 and eIF4E-3 both in the yeast two-hybrid system and FRET in *Drosophila* S2 cells.

Moreover, truncated and point mutated Me31B proteins indicate that the binding sites of Me31B for eIF4E-1 and eIF4E-3 are located in different domains. Residues Y401-L407 (at the carboxy-terminal) are essential for interaction with eIF4E-1, whereas residues F63-L70 (at the amino-terminal) are critical for interaction with eIF4E-3. Thus, Me31B represents a novel type of eIF4E-interacting protein. Our observations suggest that Me31B might recognize different eIF4E isoforms in different tissues, which could be the key to silencing specific messengers. They provide further evidence that alternative forms of eIF4E and their interactions with various partners add complexity to the control of gene expression in eukaryotes.

## INTRODUCTION

Translation is the most dynamic process to regulate the composition and quantity of a cell’s proteome. Translational control enables rapid changes in the translatability of mRNAs in response to environmental, physiological, and developmental clues (1–3). Part of this regulation occurs in large, membrane-less cumuli of assembled proteins and non-translating mRNAs termed mRNP granules or cytoplasmic foci. RNP granules form a summation of multivalent protein–protein, RNA–RNA, and protein–RNA with specific biochemical properties to define the granules (4). The different cytoplasmic foci include processing-bodies (PBs), present in most eukaryotic cells, stress granules (SGs), which form upon stress stimuli, germ granules (GGs), and neuronal granules (NGs), among others (5–7).

Messengers not being translated can be transiently stored or even degraded in some of these granules, including PBs. PBs occur in unstressed cells but can be further increased under a variety of stress conditions (5–7). PBs contain the mRNA decay machinery and, accordingly, contain enzymes involved in RNA degradation, do not present exosome components nor ribosomal proteins or translation factors (5–7). Indeed, a mutually excluding functional relationship between PBs and translation regulation has been demonstrated, as drugs that block polysomes dynamic (e.g., puromycin) promote the assembly of PBs (8–10). Intriguingly, and despite PBs contain translationally repressed mRNAs, the sole translation initiation factor present in these granules is eukaryotic translation initiation factor 4E (eIF4E). In mammals (11–14), *Xenopus* (15), *Trypanosoma brucei* (16), the planaria *Dugesia japonica* (17), *Saccharomyces cerevisiae* (18,19), *Drosophila melanogaster* (9,20), and *Caenorhabditis elegans* (21), eIF4E occurs in PBs.

eIF4E recognizes the cap structure at the 5’ end of the mRNAs. Along with the scaffold protein eIF4G and the DEAD-box ATP-dependent RNA helicase eIF4A, eIF4E conforms the eIF4F complex, which promotes cap-dependent translation initiation by mediating the interaction between the mRNA and the 43S ribosomal preinitiation complex (22,23). Evidence is mounting that eIF4E also plays critical roles in mRNA transport, storage, and translational repression in cytoplasmic foci (24). In *Xenopus* and human cells, translation of mRNAs is repressed in PBs when eIF4E interacts with eIF4E-transporter (4E-T) (11,12) and with DEAD-box ATP-dependent RNA helicase rck/p54 (humans) (12) or its *Xenopus* ortholog Xp54 (15), another component of some cytoplasmic foci, including PBs.

A key role in translation repression also has been established for the *Drosophila* ortholog of rck/p54 and Xp54, namely Maternal expression at 31B (Me31B). Along with the RNA-binding protein Tral, the mRNA 3’-UTR-binding protein Orb, the eIF4E-binding protein CUP, and the RNA localization factor Exuperantia (Exu), Me31B assembles with mRNAs to form translationally repressed mRNPs in germ granules and PBs (20,25–27). Indeed, Me31B downregulation during oogenesis results in premature translation of Oskar and Bicoid mRNAs in nurse cells (28). Recently, Wang et al. (2017) showed that Me31B is a general regulatory factor that binds to and represses the expression of thousands of maternal mRNAs during the maternal-to-zygotic transition (29). Because Me31B is expressed in different tissues throughout *Drosophila* development, i.e., nurse cells, oocytes, and early embryos, and accumulates in diverse RNP granules, such as PBs, germ plasm granules, nuage granules, and sponge bodies (20,25–28,30), Wang et al. suggested that the repressive capabilities of Me31B depend on the different biological context in which it exists (29).

PBs formation and function require protein-protein interactions and inactive mRNAs. However, an unresolved issue is how PBs can store specific mRNAs. It might involve multiple signals and proteins, some common for the whole transcriptome and some specific to a particular mRNA (31). Thus, to better understand the role of Me31B in different cell processes, it is crucial to determine the proteins interacting with it.

In this study, we analyzed the interaction in PBs of Me31B with eIF4E-1 and eIF4E-3 in *Drosophila*. We show that residues at the carboxy-terminal of Me31B (Y401-L407) are essential for interaction with eIF4E-1. In contrast, residues near the amino terminus (F63-L70) are required for the interaction with eIF4E-3, thus representing a novel structure of an eIF4E-interacting protein (4E-IPs). Our results provide further evidence to the hypothesis that alternative paralogs of eIF4E and their interactions add complexity to the control of gene expression in eukaryotic cells.

## Results

### Different eIF4E isoforms and Me31B are present in PBs in *Drosophila*

We investigated the presence of *D. melanogaster* Me31B in *S2* cells by immunofluorescence. We found Me31B in cytoplasmic foci that we considered PBs as Me31B colocalizes with GW182, a marker of PBs (Fig. 1A). In contrast, in cells stressed with sodium arsenite, we did not find that Me31B colocalizes with TIA-1, a stress granule marker (Fig. 1B). We also transfected cells with the fusion proteins YFP-Me31B or CFP-Me31B and detected the fluorescent proteins in discrete PBs, although a fraction remained dispersed in the cytoplasm (Fig. 1C).

**Figure 1.**
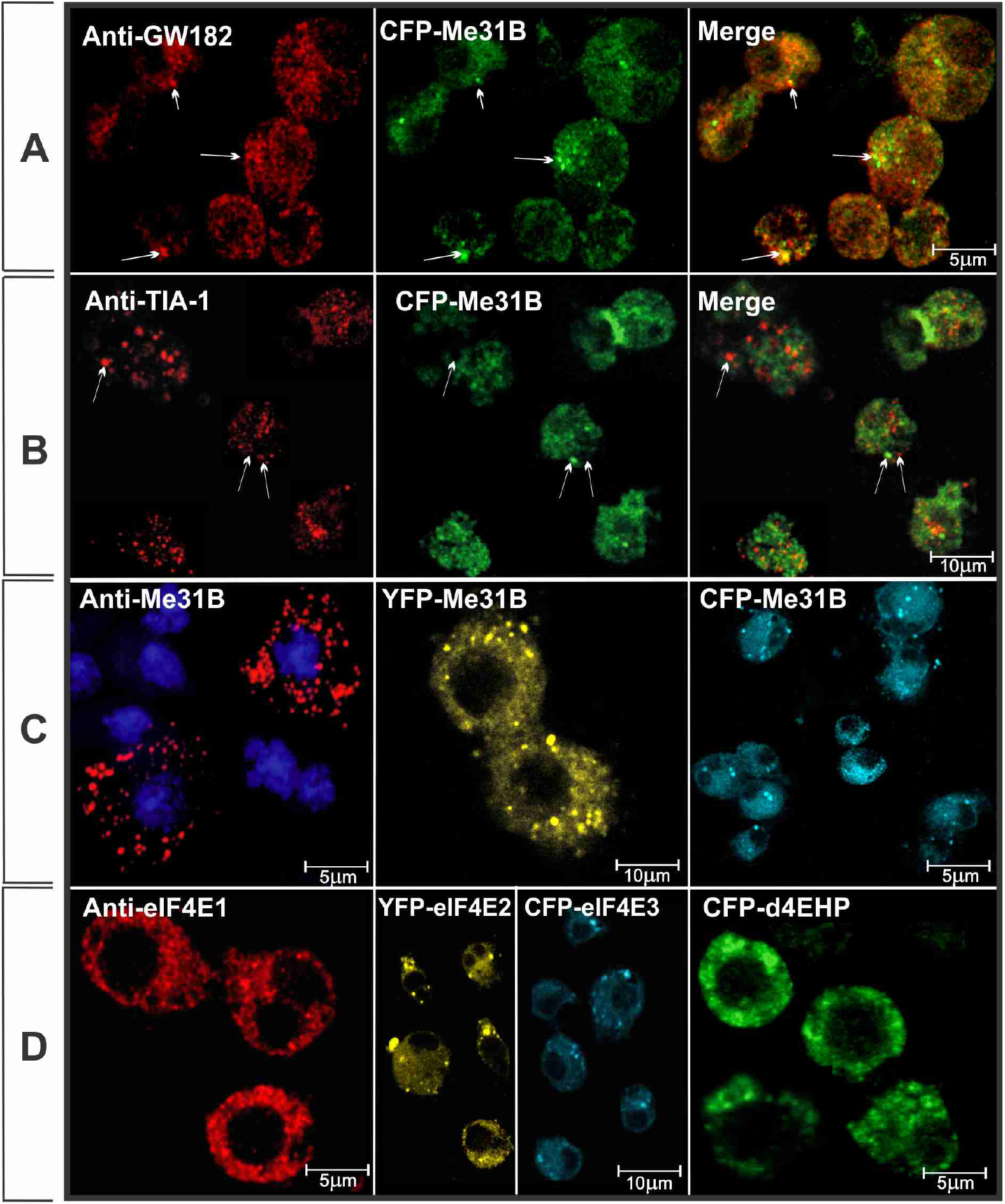
Localization of Me31B and eIF4Es in PBs. **A, B)** Me31B localizes in PBs but not stress granules. S2 cells were transfected with fluorescent protein constructs (CFP-Me31B) and immunostained with the indicated antibodies. Anti-GW182 antibody was used as PBs marker **(A)** and anti-TIA-1 antibody as a marker of stress granules **(B)**. In **(B)** cells were incubated for 30 min with 1mM sodium arsenite before fixation. Merge images show Me31B accumulation in PBs (*white arrows*). In contrast, stress granules that contain TIA-1 did not merge with CFP-Me31B (*white arrows*). **C, D)** S2 cells were either immunostained with the indicated antibodies or transfected with fluorescent protein constructs (YFP- or CFP-), as indicated. **C)** Me31B. In all cases, the indicated proteins accumulated in PBs. **D)** From left to right: eIF4E-1, eIF4E-2, eIF4E-3 and d4E-HP.

In *Drosophila* seven *eIF4E* paralog genes exist that encode eight proteins, namely eIF4E-1, eIF4E-2, eIF4E-3, eIF4E-4, eIF4E-5, eIF4E-6, eIF4E-7 and d4EHP/eIF4E-8 (32). We analized the presence of four eIF4E isoforms in *Drosophila* S2 cells either with specific antibodies, in the case of anti-eIF4E-1 or with fusion proteins in the case of YFP-eIF4E-2, YFP-eIF4E-3 and CFP-d4EHP (Fig. 1D). In all cases we detected the occurrence of eIF4E in PBs.

It has been described that eIF4E and rck/p54 colocalize in vertebrate cells (12). Therefore, we studied whether the *Drosophila* orthologous proteins colocalize in S2 cells. Simultaneously, we analyzed protein localization by cotransfection with plasmids encoding the fluorescent fusion proteins YFP-Me31B and CFP-eIF4E-3. The intensity profile across one PBs showed that colocalization is specific in cytoplasmic granules (Fig. 2). The same analysis was performed with the pair CFP-Me31B and YFP-eIF4E-1, and it was also found that both colocalize in PBs (Fig. 3). Both YFP-eIF4E-2 and YFP-d4EHP fusion proteins colocalized with CFP-Me31B as well (Fig. S1). These results suggested that endogenous Me31B interacts with eIF4Es in the same granules, which might play an important role in the PBs formation.

**Figure 2.**
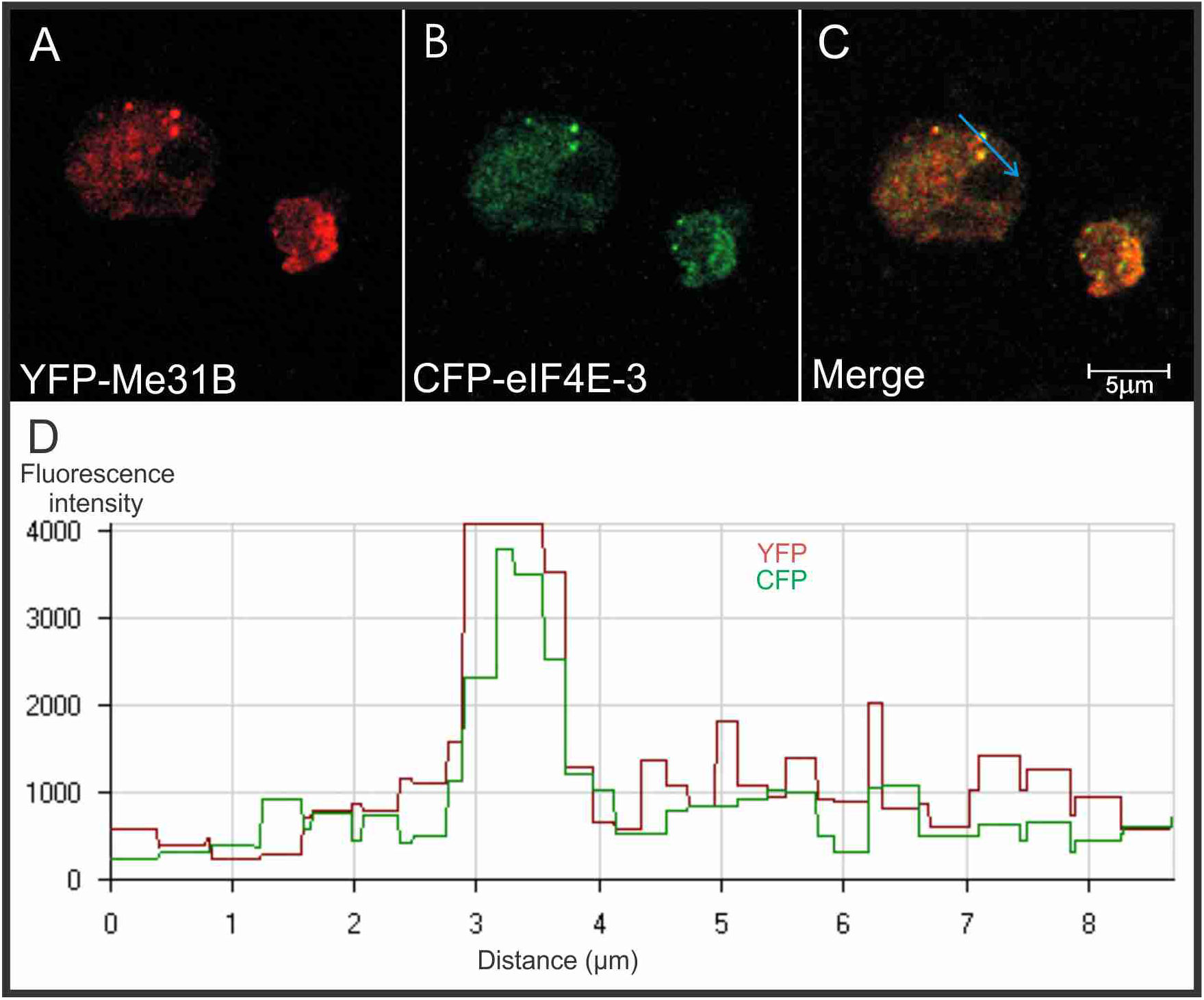
YFP-Me31B and CFP-eIF4E3 colocalization in PBs. Processing bodies of S2 cells transfected with fluorescent fusion proteins contain Me31B (A) and eIF4E-3 (B). C) In merge image we can see both proteins are located in the same granules. D) Intensity profile of CFP and YFP fluorescence as a function of the distance that a PB crosses (blue arrow in C). The superposition of both profiles confirms colocalization.

**Figure 3.**
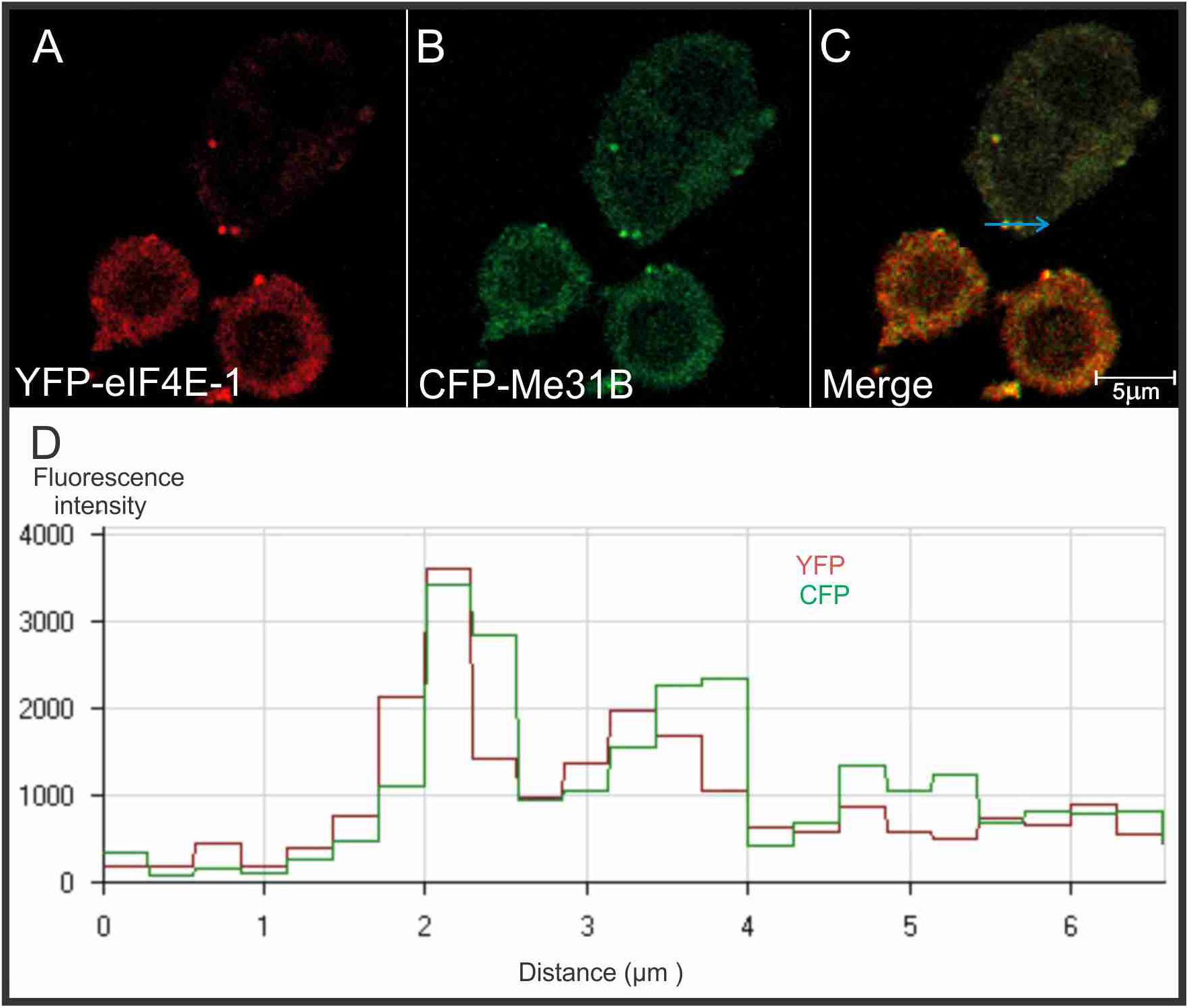
YFP-eIF4E-1 and CFP-Me31B colocalization in PBs. S2 cells transfected whith fluorescent fusion proteins contain Me31B (A) and eIF4E-1 (B). C) In merge image we can see both proteins are located in the same granules. D) Intensity profile of CFP and YFP fluorescence as a function of the distance that two PBs crosses (blue arrow in C). The superposition of both profiles confirms colocalization.

### Me31B interacts with eIF4E in PBs in *Drosophila* S2 cells

To assess protein-protein interactions, we first performed two-hybrid assays in yeast using Me31B as “prey” and eIF4E-1 and eIF4E-3 as “baits” (Fig. 4A). We observed that Me31B interacted with both eIF4E-1 and eIF4E-3. We further evaluated the protein-protein interaction in cytoplasmic granules in S2 cells using fluorescence resonance energy transfer (FRET). We tagged Me31B, eIF4E-1, and eIF4E-3 with CFP as acceptor and YFP as donor (33). FRET efficiency was measured by acceptor photobleaching, which implies that the acceptor quenches the donor fluorescence as the excitation energy is transferred to the acceptor, but after photobleaching of the acceptor, the quenching is blocked and the donor fluorescence increases. Quantification of the increase is a reliable and robust measure of FRET (34,35). Cells expressing fluorescent tagged eIF4E-1 and Me31B showed an average FRET efficiency of 36%, and cells expressing fluorescent tagged eIF4E-3 and Me31B showed an average FRET efficiency of 25%. In contrast, cells expressing only CFP and YFP (negative control) did not show significantly measured FRET (Fig. 4B). Figure 4C shows a color image of a cell coexpressing YFP-eIF4E-1 and CFP-Me31B pre and post bleached PB. In post-bleached images, we observed a diminution of the intensity of donor molecules (YFP, yellow). We show the intensity profile of CFP fluorescence before and after photobleaching, depending on the distance through the red arrow, which crosses one bleached PB. After photobleaching, CFP-fluorescence was incremented because the acceptor was absent, and the energy transfer to acceptor did not happen. The nonbleached PBs did not exhibit FRET.

**Figure 4.**
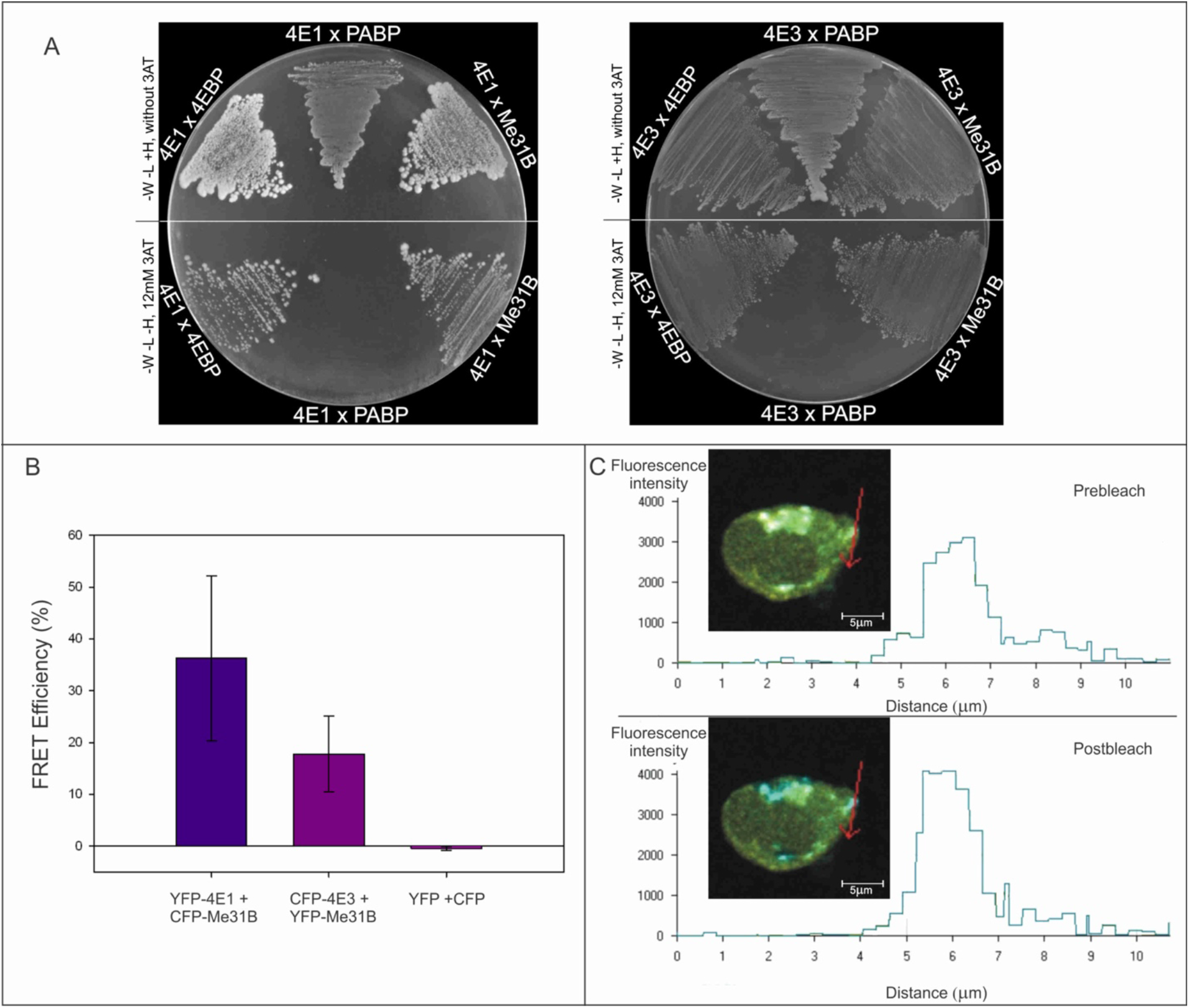
Me31B interacts with eIF4E-1 and eIF4E-3. **A)** Yeast two-hybrid system. 4E-BP was used as positive control and PABP as a negative control. *L*, leucine; *W*, tryptophan; *H*, histidine; *3AT*, 3-amino-1,2,4-triazole. **B)** FRET in S2 cells. Bar charts represent the mean values of the apparent FRET efficiencies of protein pairs from several PBs in different cells. Error bars indicate the standard deviation from the mean values. Confocal images of both CFP and YFP channels were taken before and after photo bleaching (**C**). Intensity profile of the CFP fluorescence before and afterphotobleaching as a function of the distance (*red arrow*) crossing a PB. CFP fluorescence increased after bleaching because the acceptor is absent.

The evidence that Me31B interacts with eIF4E-1 and eIF4E-3 in the yeast two-hybrid system and in S2 cells strongly suggests functional conservation with the human ortholog proteins. Therefore, we support the notion that these proteins play a role in the remodeling of mRNP.

### eIF4E residues required for PBs recruitment are essential for the interaction with Me31B

The genome of *D. melanogaster* encodes eight eIF4E isoforms (32,36). The canonical isoform, eIF4E-1, plays a role in translation and in the formation of PBs, and is a translational repressor in oogenesis when it interacts with Me31B (27,37). In eIF4E-1, the residues W100 and W146 are required for cap recognition and are involved in translation initiation, while W117 is required for protein–protein interactions and participates in translation repression and PBs assembly (9). eIF4E-3 binds to the cap structure, and both eIF4G and 4E-BP, and shares 59% identity in the carboxy-terminal moiety with eIF4E-1 (32). eIF4E-3 contains a phenylalanine residue in position 103, equivalent to W117 and W73 in *Drosophila* eIF4E-1 and human eIF4E, respectively. In figure 1D we have shown that eIF4E-1 and eIF4E-3 localized in cytoplasmic granules, and previously was demonstrated that the mutant eIF4E-3^F103A^, like eIF4E-1^W117A^, does not localize in cytoplasmic granules (9). Yeast two-hybrid experiments showed that eIF4E-1 W117 is required to interact with Me31B (Fig. 5). Similarly, eIF4E-3 / Me31B interaction required F103 residue. These results suggest that the interaction between Me31B and eIF4E-1 and −3 might be required to recruit of eIF4Es to PBs, most possible for silencing mRNAs.

**Figure 5.**
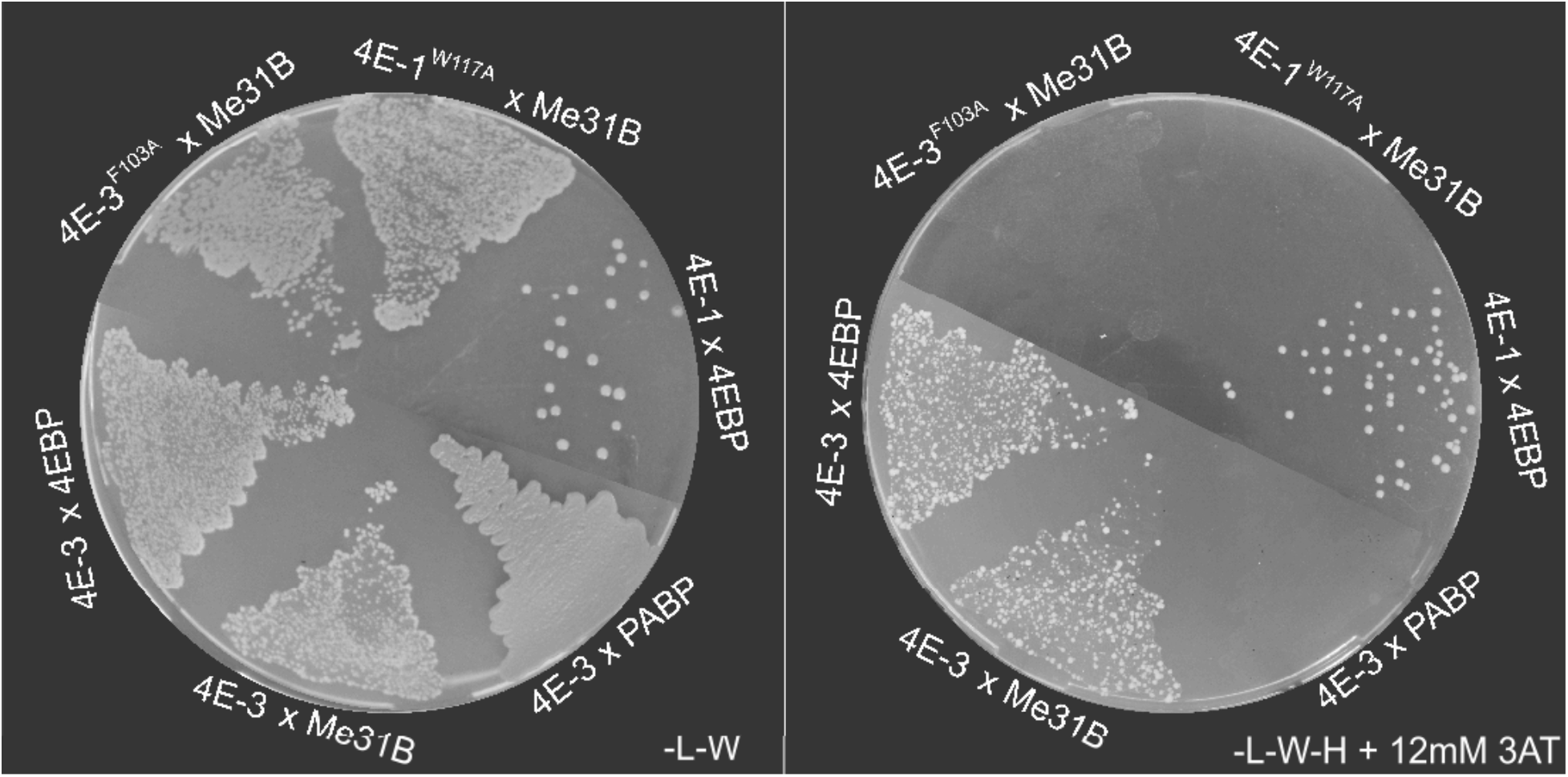
Mutants of eIF4E at essential sites for accumulation in PBs do not interact with Me31B. eIF4E-1^W117A^ and eIF4E-3^F103A^ do not interact with Me31B in the yeast two-hybrid system. *Left:* mating control plate. 4E-BP was used as positive control and PABP as a negative control. *L*, leucine; *W*, tryptophan; *H*, histidine; *3AT*, 3-amino-1,2,4-triazole.

### Different domains of Me31B interact with eIF4E-1 and eIF4E-3

Me31B is a member of a highly conserved superfamily 2/DDX6 of DEAD-box RNA helicases with roles in translational repression from trypanosomes to humans (38,39). Me31B is composed of two linked RecA-like domains, domains 1 and 2. Domain 1 contains the Q motif followed by motifs I–III, and domain 2 contains motifs IV–VI participate in ATP binding and RNA binding (Fig. 6D). As eIF4E-1 and eIF4E-3 coexist in different tissues, we asked whether the interaction is mediated by the same or different domains of Me31B to determine competitive or simultaneous interactions.

**Figure 6.**
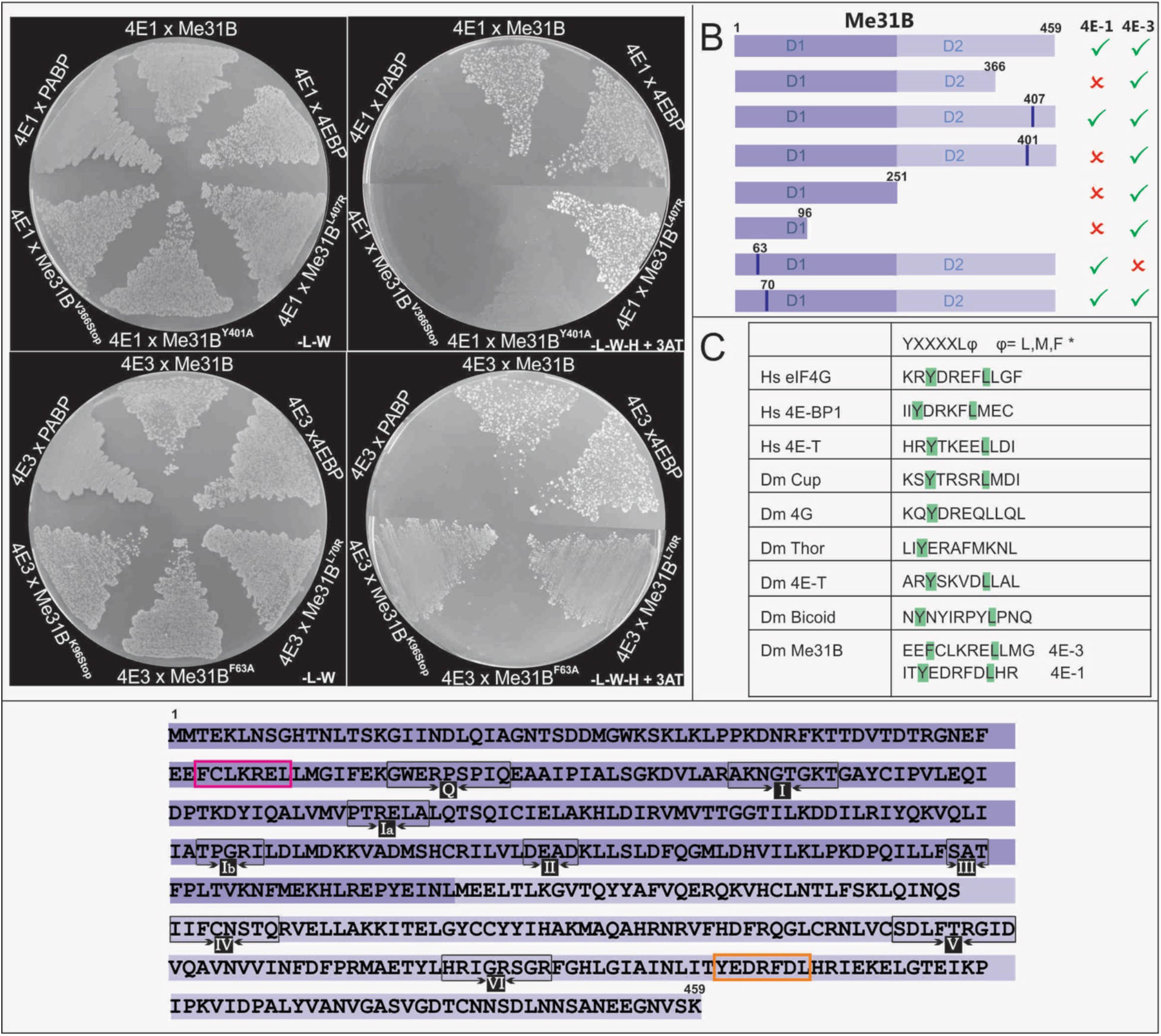
Two independent domains of Me31B interact with eIF4E-1 and eIF4E-3. **A)** Yeast two-hybrid assays with Me31B mutants. *Left*, mating control plate. 4E-BP was used as positive control and PABP as a negative control. *L*, leucine; *W*, tryptophan; *H*, histidine; *3AT*, 3-amino-1,2,4-triazole. **B)** Me31B protein fragments are represented as *bars*. D1 and D2 domains are highlighted in *dark purple* and light purple, respectively. Different mutants were evaluated for interaction. *Crosses* indicate no interaction, and *ticks* indicate positive interactions. **C)** Consensus sequences of the eIF4E-binding motif (4E-BM) from the indicated eIF4E-interaction proteins. Me31B 4E-BMs found in this study are also shown. **D)** Me31B sequence. Interaction sites with eIF4E-1 and eIF4E-3 are marked in *orange* and *pink*, respectively.

We used the yeast two-hybrid system assays to analyze the interaction of truncated Me31B with both eIF4Es. We mutated the codon of Valine 366, Cysteine 285, Serine 251, Lysine 194, Lysine 124, and Lysine 96 to stop codon (Fig. 6B). Our results indicated that the binding sites of Me31B for eIF4E-3 and eIF4E-1 are located in different domains of Me31B at the amino and carboxy-terminus, respectively (Fig. 6A-B). eIF4E-3 bound to D1 domain (Me31B^1-96^) whereas eIF4E-1 bound to D2 domain (Me31B^366-459^). Next, we wanted to identify the two eIF4E-binding sites (4E-BSs) present in Me31B. A comparison of the 4E-BSs occurring in other eIF4E-interacting proteins suggested two putative 4E-BSs within the regions identified of Me31B (Fig. 6C). To test the functionality of the putative 4E-BSs, we performed site-directed mutagenesis to replace residues F63 to Ala and L70 to Arg in the eIF4E-3 interacting region and Y401 to Ala and L407 to Arg in the eIF4E-1 interacting site. The substitution of both aromatic residues Y401 and F63 to a non-aromatic residue prevented the interaction of Me31B with eIF4E-1 and eIF4E-3, respectively (Fig. 6A-B). Therefore, we conclude that Me31B interacts with eIF4E-1 and eIF4E-3 through independent binding sites specific for each isoform. Thus, Me31B represents a novel eIF4E-interacting protein.

### Expression of Me31B and eIF4E-3 in the male germ line

The independent binding of the eIF4E isoforms with Me31B led us to consider the biological significance of this remarkable feature. eIF4E-1 is ubiquitously expressed in *D. melanogaster*, while the function of eIF4E-3 is restricted to spermatogenesis (40). Therefore, we analyzed whether Me31B is expressed in the male germline. We first assessed the presence of the Me31B messenger in testis cells (Fig. S2), and then we analyzed the location of Me31B by means of a transgenic line expressing a GFP-ME31B fusion. We detected Me31B-GFP protein in cytoplasmic granules in a subset of male germ cells (Fig. 7A). Moreover, we observed the co-expression of Me31B with eIF4E-3 (Fig. 7B-C). These results suggest a biological function of Me31B in the male germline. As the spermatogenesis proceed, we identified Me31B and eIF4E-3 associated to the meiotic chromosomes (Fig. 7D-F). This agrees with the described function of eIF4E-3 in chromosome segreggation and opens a path for further studies on the biological of Me31B in the male germ line.

**Figure 7.**
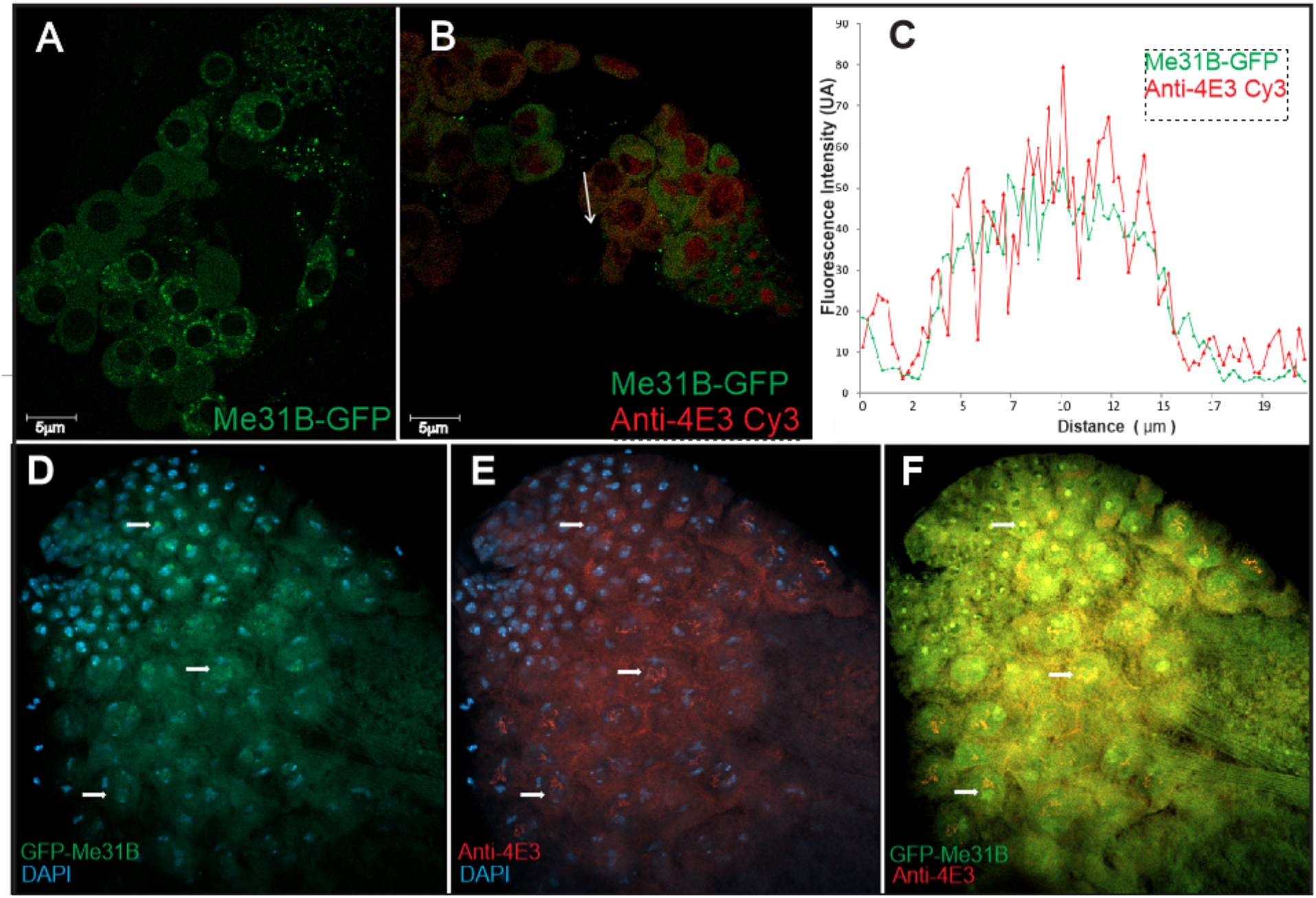
Me31B is found in testis cells and colocalizes with eIF4E-3. **A)** Me31B is observed located in cytoplasmic granules in testis cells of the trap line GFP-Me31B flies. **B)** Testis immunofluorescence with anti-eIF4E-3 antibodies of GFP-Me31B flies. Both proteins are located in the same cells **(C)**. Intensity profile of GFP and Cy3 fluorescence as a function of the distance that a PB crosses (*white arrow* in **B**). The superposition of both profiles confirms colocalization. **7D, E)** Me31B and eIF4E-3 are seen associated with meiotic chromosomes, DAPI staining is showed in blue. The colocation of both is shown in figure **(F)**.

## Discussion

The presence of several paralogs of eIF4E in *Drosophila* (32,36) and other species led to the hypothesis that eIF4E is the sole translation factor present in active mRNAs and in the mRNP of PBs (41). Thus, we could hypothesize that Me31B is involved in the transition from active mRNA to a state of repression in PBs and it could play different roles in assemblying distinct mRNP complexes. An ubiquitous set of eIF4 factors might support global and basal translation initiation, whereas other isoforms might regulate the translation of a subset of mRNAs in a tissue- and or developmental stage-specific manner (41). *Drosophila* eIF4E-1 seems to be the canonical isoform (32), while eIF4E-8/d4E-HP is a repressor of *bicoid* mRNA during embryogenesis (42), and eIF4E-3 is a testis-specific modulator of male germline development (40,43).

Cytoplasmic mRNA granules can be found in a variety of configurations, depending on their protein content. These structures play critical roles in post-transcriptional regulation of gene expression (31,44). The interaction of *Drosophila* eIF4E-1 and eIF4E-3 with Me31B in PBs agrees with RNAi studies demonstrating that Me31B is necessary for PBs assembly (45). Me31B also plays an essential role in translation regulation during oogénesis (28). Thus, Me31B along with other proteins, might be involved in mRNPs remodeling from active polysomes to repression by direct interaction with eIF4E. A model has been previously proposed by the removal of the translation machinery, mRNP reorganization, and PBs assembly (31). Subsequently, other PBs components would be recruited for mRNA degradation (Dcps, Xrn1 enzymes, among others). The evidence shown here that the mutants eIF4E-1^W117A^ and eIF4E-3^F103A^, which are prevented from being assembled into PBs (9), do not interact with Me31B supports this notion. A plausible mechanism might be that when Me31B does not bind eIF4E, it cannot remodel the active mRNPs for silencing into PBs.

We also have shown that Me31B can interact with different eIF4E isoforms through different 4E-BS. While the eIF4E-3-binding site is located at the N-terminus, eIF4E-1 interaction is mediated by a 4E-BS in the C-terminus. We have also shown that both sites are isoform-specific, a remarkable feature of Me31B. These unique features render *Drosophila* Me31B the first protein so-far reported with such capabilities. These results agree with the idea suggesting that the repressive capacities of Me31B depend on the various interactors and different biological contexts in which it exists (29). Both Me31B 4E-BSs are non-canonical motifs and might interact with different eIF4E domains, as it was shown for the d4E-HP/Bicoid interaction (42). Recent studies showed that some 4E-BPs need two domains to interact with the same eIF4E, namely one canonical 4E-BS that interacts with the dorsal surface of eIF4E, and a non-canonical one that interacts with its lateral surface (46). Other 4E-BPs, such as *Drosophila* Mextil, uses a third interaction motif (auxiliary motif) to interact with eIF4E-1 (47). Our data add up to the complexity of the interacting mRNP landscape showing isoform-specific independent binding sites within the same eIF4E-interacting protein.

Our current working hypothesis is that Me31B plays a crucial role in mRNP remodeling and mRNA silencing in the female and male germline, by interaction with eIF4E-1 in the ovaries or by interaction with eIF4E-3 in testes.

Previous work had demonstrated that *Drosophila* eIF4E-3 plays a critical role in spermatogenesis (40,43). However, while 4E-BP binds eIF4E-1 in testes, it does not bind eIF4E-3, raising the critical question of what protein or proteins regulate eIF4E-3 during this developmental process (40,43). Our results here strongly support the notion that Me31B might be the eIF4E-interacting protein that controls eIF4E-3 activity throughout male germline development. Further analysis of the function and structure of the Me31B/eIF4E-3 interaction in testes will shed light on the appealing hypothesis of a common germline regulation mechanism.

The results described here and the previous evidence from us and other laboratories further support the notion of the plasticity for the eIF4E interaction network, which confers unique properties to the different assembled mRNPs and gene expression control in eukaryotes.

## EXPERIMENTAL PROCEDURES

### Cell culture and transfections

*Drosophila* S2 cells were grown on glass coverslips (Fisher Scientific) in Schneider’s medium (Sigma, USA), supplemented with 10% fetal bovine serum (Natocor, Córdoba, Argentina) and 1% antibiotic–antimycotic mixture (Invitrogen, USA) at 28 °C. Plasmid transfections were performed after cells had reached ~90% confluency using Lipofectamine reagent (Roche), as recommended by the manufacturer. Sixteen hours after transfection, cells were washed with PBS (130mM NaCl, 20 mM potassium phosphate, pH 7.4), fixed for 20 min with PBS pH 7.4/4% w/v paraformaldehyde, and mounted in antifade (Mowiol, Calbiochem).

### Immunofluorescence

Cells were fixed as described above, washed with PBS pH 7.4, and permeabilized in PBS pH 7.4/0.2% Triton X-100 (Sigma) for 20 min. Cells were then rinsed with PBS, blocked in PBS pH 7.4 / 10% fetal calf serum (FCS) for 30 min, and incubated with the primary antibody diluted in PBS pH 7.4/10% FCS for 60 min. Subsequently, cells were washed with PBS pH 7.4 (4 × 15 min) and incubated with the secondary antibody in PBS pH 7.4 / 10%FCS for 45 min. Cells were again washed with PBS pH 7.4 (3 × 10 min) and mounted in antifade (Mowiol, Calbiochem).

### Confocal laser scanning microscopy

Before imaging, cells were counterstained with DAPI and analyzed by epifluorescence to assess cell integrity. Images were acquired with a Carl Zeiss LSM 510-Meta confocal microscope using Argon (588/514 nm) and Helium/Neon (543/633 nm) lasers. The images were analyzed using the LSM software and Image J (http://rsbweb.nih.gov/ij/).

The following primary antibodies were used in this study: rabbit anti-rck/p54 (DDX6, Bethyl Laboratories; 1:500); rabbit anti-eIF4E-1 1:1000 (48); rabbit polyclonal affinity-purified anti-eIF4E-3 antibodies #967 and #968 1:300 (Biomatik, Ontario, Canada, (40)) anti-GW182 and anti-TIA-1 (AbCam, Cambridge, UK; 1:500). The following secondary antibodies were used: anti-mouse, anti-goat, and anti-rabbit antibodies conjugated to Cyanine dyes (Jackson Inc.; 1:2000).

### Acceptor photobleaching FRET

All data were obtained on an LSM 510 Meta (Zeiss). Samples were fixed with 4% PFA for 15 min and images acquired with a CAPOCHROMAT 63×/1.4 W Korr objective (Zeiss). Specific excitation and emission of the CFP-fusion proteins were effected by excitation at 458 nm with a 30 mW Argon/2 laser (AOTF transmission 15%) and collection of emitted light with a 475/525 nm bandpass filter. No emission from YFP fusion proteins was detected in this channel. CFP images were taken before and after photobleaching of the YFP signal using the same sensitivity settings. YFP signals were photobleached by full-power excitation at 514 nm with a 50-mW solid-state laser. Images of YFP-fusion proteins were obtained before and after photobleaching by excitation with a 30-mW Argon/2 laser (transmission 15%) at 514 nm excitation and emission collected from 530 nm bandpass filter (LSM 510 Meta Detector, Zeiss). No photobleaching of the CFP signal was observed under >90% photodestruction of the YFP signal. The FRET efficiency (49) was determined for each PB separately by: Eap = (ID0-ID)/ ID0; with ID0 and ID denoting the sum of the respective pre- and post bleach donor intensity in a PB.

### Immunohistochemistry of Trap-line GFP-Me31B flies

*y1-w1118; P{PTT-GB}hme31* BCB05282 fly stock was obtained from the Bloomington *Drosophila* Stock Center. Testes were dissected on ice in S2 Schneider’s medium supplemented with 10% fetal bovine serum. The medium was removed and testes were fixed in fixer solution [200 μl of 4% paraformaldehyde in PBST (PBS with 0.2%Tween 20), 600 μl heptane and 20 μl DMSO] for 20 minutes with slow rotation. The fixer was removed, and testes were washed three times for 15minutes each in 1.5 ml PBST followed by 1-2 hours blocking with 1 ml blocking solution (PBST, 0.1% Triton X-100, 1% BSA). Testes were incubated either with primary antibodies in blocking solution at 4°C overnight, washed for 30 minutes in PBST, blocked for 30 minutes with 500 μl blocking solution containing 1% goat serum, and incubated with secondary antibodies in 500 μl blocking solution containing 8% goat serum overnight at 4°C. At this step, testes were counterstained with DAPI (1 ng/ml) for 5 minutes, washed four times for 15 minutes each with 1.5 ml PBST, and mounted for imaging.

### Plasmids construction

We generated plasmids encoding the YFP and CFP fusion proteins. *Drosophila* eIF4E-1 (FlyBase CG4035), eIF4E-2 (FlyBase CG4035), eIF4E-3 (FlyBase CG 8023), d4E-HP (FlyBase CG33100) (32,36) and Me31B (FlyBase CG4916) (28). cDNAs were PCR-amplified and cloned as EcoRI-EcoRI fragments onto the Topo Blunt vector (Invitrogen). cDNAs were PCR-re-amplified and finally subcloned into Hind III site of the pEYFP-C1 or pECFP-C1 vectors (Clontech).

*Drosophila* eIF4E-1 and eIF4E-3 cDNAs (32) were cloned into the pOBD2 vector (50) in-frame with the DNA-binding domain sequence of GAL4 to create the “bait” constructs. Me31B, 4E-BP (CG8846; (51)) and PABP (CG5119; (52)) were cloned into the activator domain sequence of GAL4 to generate the “prey” constructs. Me31B and 4E-BP were cloned into pGAD424 vector and PABP into pACT2 vector. See table S1 for a complete list of the plasmids used in this work.

### Yeast Two Hybrid Assay

Interactions between “bait” and “prey” proteins were detected following a yeast interaction-mating method using the strains PJ69-4a and PJ69-4alpha (table S2; (50)). Diploid cells containing both bait and prey plasmids were grown in selective media (–Trp, –Leu) and showed as growth control. Protein interactions were detected by replica-plating diploid cells onto selective media –(Trp, Leu, His) containing 3-amino-1,2,4-triazole (3AT). Growth was scored after four days of growth at 30°C.

The 3AT amount was titled within a range of 0, 10, 12, 15, 20, and 30mM of 3AT, resulting in 12mM the minimum concentration needed to inhibit background growth and still enable the positive growth’s controls.

### Site direct mutagenesis

Site-directed mutagenesis of eIF4E-1 was carried out on the plasmids pOBD-eIF4E1 to change tryptophan 117 to alanine, and pOBD-eIF4E-3 was used to change Phenylalanine 103 to alanine. PCR amplification of each template was performed using 2X iProof master mix (Bio-Rad) with primers described in Ferrero *et al* 2012. The same protocol was used to mutate pGAD424-Me31B with the following primers (for more details, see Table S3):

Me31B^F63A^r: CTTTTAAGGCAAGCCTCCTCGAA
Me31B^F63A^f: TTCGAGGAGGCTTGCCTTAAAAG
Me31B^L70R^r: GAATATACCCATACGCAGTTCTC
Me31B^L70R^f: GAGAACTGCGTATGGGTATATTC
Me31B^V366STOP^r: AATTGATTACTTAATTCACGGC
Me31B^V366STOP^f: GCCGTGAATTAAGTAATCAATT
Me31B^K251STOP^r: GCGTAAATGTTACTCCATGAA
Me31B^K251STOP^f: TTCATGGAGTAACATTTACGC
Me31B^Y401A^f: CTGATAACCGCCGAGGATCGGTT
Me31B^Y401A^r: AACCGATCCTCGGCGGTTATCA
Me31B^L407R^f: CGGTTTGATCGGCATCGGATTGA
Me31B^L407R^r: TCAATCCGATGCCGATCAAACCG

## ACKNOWLEDGEMENTS

We thank Paul Lasko (McGill University, Canada) for *Drosophila* anti-eIF4E-3 antibodies.

## AUTHOR CONTRIBUTIONS

Conceived and designed the experiments: C.L. and R.R.P. Performed the experiments: G.C., E.V. and C.L. Analyzed the data: C.L., R.R.P and G.H. Contributed reagents/materials/analysis tools: C.L. and R.R.P. Wrote the paper: G.H., R.R.P. and C.L.

## FUNDING AND ADDITIONAL INFORMATION

This work was supported by grants of ANPCyT (PICT-2016-2045 to C.L. and PICT-2015-2128 to R.R.P); C.L. and R.R.P. are investigators and E.V. is a doctoral fellow of CONICET, Argentina. G.H. was supported by the intramural funding program of the National Institute of Cancer (Instituto Nacional de Cancerología, INCan), Mexico.

